# Analysis of the *Aedes albopictus* C6/36 genome provides insight into cell line adaptations to *in vitro* viral propagation

**DOI:** 10.1101/157081

**Authors:** Jason R Miller, Sergey Koren, Kari A Dilley, Vinita Puri, David M Brown, Derek M Harkins, Françoise Thibaud-Nissen, Benjamin Rosen, Xiao-Guang Chen, Zhijian Tu, Igor V Sharakhov, Maria V Sharakhova, Robert Sebra, Timothy B Stockwell, Nicholas H Bergman, Granger G Sutton, Adam M Phillippy, Peter M Piermarini, Reed S Shabman

## Abstract

**Background:** The 50-year old *Aedes albopictus* C6/36 cell line is a resource for the detection, amplification, and analysis of mosquito-borne viruses including Zika, dengue, and chikungunya. The cell line is derived from an unknown number of larvae from an unspecified strain of *Aedes albopictus* mosquitoes. Toward improved utility of the cell line for research in virus transmission, we present an annotated assembly of the C6/36 genome.

**Results:** The C6/36 genome assembly has the largest contig N50 (3.3 Mbp) of any mosquito assembly, presents the sequences of both haplotypes for most of the diploid genome, reveals independent null mutations in both alleles of the Dicer locus, and indicates a male-specific genome. Gene annotation was computed with publicly available mosquito transcript sequences. Gene expression data from cell line RNA sequence identified enrichment of growth-related pathways and conspicuous deficiency in aquaporins and inward rectifier K^+^ channels. As a test of utility, RNA sequence data from Zika-infected cells was mapped to the C6/36 genome and transcriptome assemblies. Host subtraction reduced the data set by 89%, enabling faster characterization of non-host reads.

**Conclusions:** The C6/36 genome sequence and annotation should enable additional uses of the cell line to study arbovirus vector interactions and interventions aimed at restricting the spread of human disease.

## BACKGROUND

Insect cell lines such as Aag2 and C6/36 are critical platforms for insect biology and virology. The *Aedes albopictus* clone C6/36 (ATCC CRL-1660) cell line is commonly used for detection, propagation, and analysis of arboviruses, including antibody-based detection of viruses in saliva [reviewed in [1]]. C6/36 cells have a short population doubling time and are permissive to infection by mosquito-transmitted viruses across members of the *Togaviridae, Flaviviridae*, and *Bunyaviridae* families. In particular, C6/36 cells are used to study viruses that pose significant threats to human health, including Zika, dengue, chikungunya. Virus propagation in C6/36 cells guides the rational development of vaccines and therapeutics. PubMed [2] lists 671 publications with C6/36 in the title or abstract.

The progenitor of the C6/36 cell line was established in 1967 from freshly hatched *Aedes albopictus* larvae of unspecified ancestry [3]. The C6/36 subclone was selected for its uniformly high virus yield and was shown to retain a diploid karyotype with 2n=6 chromosomes in a majority of cells [4]. The similar or equivalent ATC-15 cells [1] were shown to be diploid [5] and to have more chromosomal abnormalities after 110 passages than after 17 [6]. The C6/36 cell line, available through the American Type Culture Collection (ATCC, Manassas VA) is described as maintaining a diploid chromosome number and being non-anchorage dependent and non-tumorigenic [7]. Despite the widespread use of this cell line to both propagate arboviruses and to use them as a tool to study virus-mosquito interactions, little has been published about features that differentiate the cell line genome from that of *Aedes albopictus* mosquitoes.

Two strains of *A. albopictus* have published genomes, both of which were sequenced on Illumina platforms and assembled with the SOAPdenovo assembler [8]. Sequencing of the Italian Fellini aka Rimini strain yielded small contigs with N50 < 1 Kbp [9]. The assembly of a Foshan female from China [10] as provided in VectorBase [11] version AaloF1, has a 1.92 Gbp scaffold span, 1.78 Gbp contig span, and 18.4 Kbp contig N50. A third strain was analyzed for its genomic repeats using a pipeline called dnaPipeTE that runs on Illumina reads [12]. The *A. aegypti* Liverpool genome was assembled to draft status from Sanger reads [13] and later de-duplicated (removing putative redundant contigs) and extended to chromosome-length scaffolds with Hi-C technology [14]. The 2014 update in VectorBase has an 82 Kbp contig N50. Using these assemblies, the within-genus divergence between *A. albopictus* and *A. aegypti* was estimated at 71.4 mya [10]. High population heterozygosity has been recognized in mosquitoes for over 35 years [15], indicating the C6/36 cells could harbor a heterozygous genome.

Recent advances in DNA sequencing technology have enabled the generation of megabase-scale contigs. The Pacific BioSciences (PacBio, Menlo Park CA) and Oxford Nanopore (Oxford UK) single-molecule sequencing platforms can generate reads in excess of 10 Kbp. Due to its randomness, the high base call error in PacBio reads can be overcome by using sequencing depths in the 50X range [16]. New assembly algorithms targeting deep-coverage PacBio data have separated the haplotypes from heterozygous regions of diploid genomes [17].

Here we describe findings for the *Aedes albopictus* C6/36 cell line including its karyotype, the assembly of its genome from PacBio sequence, an analysis of the haplotype separation in contigs, the gene annotation based on public mosquito RNA sequence, and analysis of gene expression based on cell line RNA. We also demonstrate use of the genome and transcriptome for the purpose of subtracting host sequence during an RNA sequencing assay for viruses. The sequence data were previously deposited in public databases to facilitate research on Zika and other viruses commonly transmitted by *Aedes albopictus* and studied in C6/36 cell lines.

## DATA DESCRIPTION

The assembly of the genome sequence is available at NCBI [18] under accession GCA_001876365.2. The contig accessions start at MNAF02000000. The annotation describing the transcripts and genes is available at NCBI under Aedes albopictus Annotation Release 101. The sequencing reads used to generate, validate, and analyze the assembly are available at NCBI SRA; see **Table S1 and S2** for accessions. The assembly and annotation are also available at VectorBase [11] under the name canu_80X_arrow2.2.

## ANALYSIS

C6/36 cells, obtained from ATCC (CRL-1660) and cultured at JCVI, were subjected to visual analysis of stained metaphase chromosomes to ascertain the karyotype. All cells examined displayed the three metacentric chromosomes expected of mosquito cells. A majority of cells also displayed additional chromosomes, as shown in Figure 1, and the specific composition varied per cell. This analysis suggested variable and cell-specific partial duplications of chromosomes. Notably, a whole-genome duplication was not indicated. overlap. There are several additional short metacentric and acrocentric chromosomes shown by arrows. The 5 μm scale bar applies to all three images.

**Figure 1.**
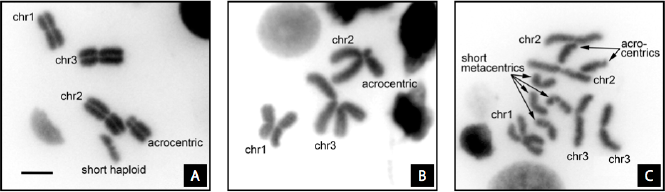
Three karyotypes of the C6/36 *Ae. albopictus* cell line. Chromosomes are labeled chr1 for the shortest, chr2 for the longest, and chr3 for the intermediate size chromosomes within each image. 1a: Cell in prometaphase has three normal paired chromosomes. An additional acrocentric chromosome pair has a short arm indicating deletion or translocation to elsewhere. An additional short haploid chromosome is unpaired. 1b: Cell in early metaphase has chromosomes with pairs slightly separated. Chr1 appears normal. The other chromosomes are abnormal, possibly due to translocation of most of one arm of chr2 to one arm of chr3. 1c: Cell in mid-metaphase with chromosome pairs separated. Chr1, chr2, and chr3 appear normal. The chr1 homologous pairs overlap. There are several additional short metacentric and acrocentric chromosomes shown by arrows. The 5 μm scale bar applies to all three images.

### Sequencing and assembly generate large contigs

Genomic long-read sequencing for assembly generated 161 Gbp in 17.9M total reads providing 147 Gbp in 12.4 M reads 5 Kbp or longer and 107 Gbp in 7.10 M reads 10 Kbp or longer. See **Table S1** for accessions. Genomic short-read sequencing for analysis generated 45.8 Gbp in 152 M pairs of 2x150bp reads. Transcript sequencing yielded 16.6 Gbp in 27.7 M pairs of 2x300bp partially overlapping reads. See **Table S2** for accessions.

Ten candidate assemblies were generated by combinations of four software packages: either the Falcon or the Canu assembler [17, 19], followed by zero to two iterations of either the Quiver or Arrow consensus polisher [20]. As shown in **Table S3**, the ten resulting assemblies had similar size profiles. The sum of bases was 2.25 Gbp in all assemblies. The average long-read coverage was 72X. Each assembly mapped 93% of the 20X paired short reads, which had not been used during assembly; see **Table S4**. Local alignments aligned the assemblies’ entire spans in large segments. For example, the contigs of Falcon and Canu after 2 rounds of Arrow each, had 2.24 Gbp in alignments of at least 99% sequence identity, and these aligned spans had 630 Kbp N50.

The assembly chosen for downstream analysis was the one from Canu plus two rounds of Arrow because its 93.2% short-read map rate was highest by a small margin. This assembly had a total span of 2.247 Gbp in 2,434 contigs and a contig N50 of 3.304 Mbp; see **Table S5**. This assembly is available in GenBank with accession GCA_001876365.2. To our knowledge, the C6/36 contig N50 exceeds that of any other assembly of any mosquito or mosquito cell line genome, though some other assemblies offer scaffolds and chromosome mappings in addition to contigs; see **Table S6**.

### Dissimilarity with other mosquito assemblies

The C6/36 assembly was compared to *Aedes albopictus* Foshan [10]. The C6/36 contig span is 28% larger. Global alignments spanned 816 Mbp (stringent) or 1.44 Gbp (permissive) of both assemblies; see **Table S7**. Local alignments covered 692 Mbp of Foshan contigs and 1,028 Mbp of C6/36 contigs; see **Table S8**. Thus, both alignment methods left large portions of both assemblies unaligned. Sequence identity within alignments was low. The local alignments with at least 95% sequence identity covered only 373 Mbp of Foshan and 596 Mbp of C6/36. Local alignments covered more of C6/36 than Foshan, indicating that some sequences are present at higher multiplicity in the C6/36 assembly. To explore the Foshan vs. C6/36 genome difference free of the C6/36 assembly, the C6/36 short read pairs were mapped to Foshan contigs. As shown in **Table S9**, 93% of pairs had mapped to C6/36 contigs but only 49% mapped to Foshan contigs. Of pairs not mapped to C6/36, less than 1% mapped to Foshan. These results combine to indicate dissimilarity of the Foshan and C6/36 genomes. It is possible that the cell line is derived from an *A. albopictus* strain that was itself diverged from Foshan. Inter-strain genome size differences had been noted in this species prior to the sequencing era [21].

Inter-species nucleotide alignment was unproductive. Permissive global alignments to *Aedes aegypti* [13, 14] covered only 15% (290 Mbp) of Foshan and 14% (304 Mbp) of C6/36.

C6/36 repeats were detected, characterized, and mapped back to the assembly with the process used for Foshan [10]. As shown in **Table S10**, results were similar to those reported for Foshan. Repeats cover 74% of C6/36 (and 76% of Foshan) and the three most abundant repeat types accounted for 59% of assembled bases (and 60% of Foshan bases). The most abundant repeat types were Unknown, LINE retrotransposon, and DNA transposon in C6/36 (and LINE, LTR retrotransposon, and Other in Foshan).

### Redundancy indicates haplotype separation

The Canu assembler can separate haplotype regions having over 2% divergence [19]. Therefore, we evaluated the C6/36 assembly for haplotype separation. The first evaluation used the genomic short reads, which were not used by the assembler. K-mer analysis of the short reads, independent of the assembly, estimated 5.7% heterozygosity within the C6/36 genome and a genome size less than half the assembled contig span; see **Figure S1a**. Mapping the reads to contigs yielded an overall coverage mode of 18X; see **Figure S1b**. The coverage mode for most individual contigs was also about 18X; see Figure 2 and **Table S11**. The bimodal coverage is similar to that seen in the *Quercus lobata* (oak) genome assembly, Figure 2B in [22], for which the assembler putatively separated the haplotypes at divergent loci. In C6/36, the 1832 contigs whose mode coverage was in the 18X ± 6X range collectively span 95.82% of the assembled bases. There is a smaller group of large contigs with mode coverage in the 36X ± 6X range. These results support the hypothesis that most of the C6/36 diploid genome is represented twice in the assembly, possibly due to haplotype separation at heterozygous loci, such that collapsed sequences from less heterozygous loci attracted twice as much short-read coverage.

**Figure 2.**
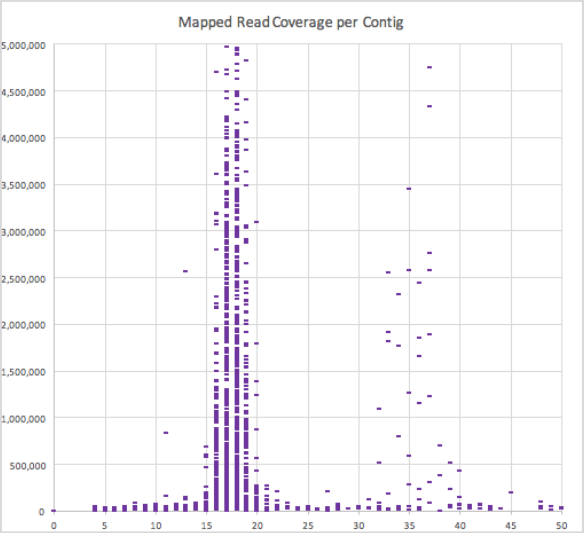
Contig size vs coverage. Each bar in the scatter plot represents one contig to which short reads were mapped. Y-axis: contig size in bases. X-axis: mode of the short-read coverage of one contig, in bins of 1X. The 1832 contigs with mode coverage in the 18X ± 6X range collectively span 95.82% of the assembled bases, but there are a few large contigs with 36X ± 6X coverage. The apparent bi-modal distribution suggests that the ~18X contigs could contain separate representations of heterozygous loci, while the ~36X contigs could contain the consensus of both alleles from less heterozygous loci.

Second, the C6/36 assembly was tested for the presence of sequence-similar contigs. After alignment of the nucleotide sequence to itself was inconclusive, short reads were mapped allowing up to 4 maps per pair, and the 15% of read pairs that mapped exactly twice were used to identify paired contigs (PCs), defined here as two contigs sharing at least 10,000 read pairs that mapped twice. This identified the 529 PCs described in **Table S12**. There were 708 contigs in PCs (474 contigs in exactly one PC and maximum 8 PCs for one contig). There were 689 PC contigs with short-read coverage modes in the 18X ± 6X range (maximum coverage was 51X). The PCs incorporated 1.97 Gbp or 88% of the 2.25 Gbp assembly. Sequence similarity within PCs was low; average 93.5% identity in aligned bases, with 28% of the sequence aligned (274 Mbp in alignments covering 548 Mbp on contigs). Where PC alignments left one contig extending past the other, it was possible to “walk” from PC to PC as illustrated in **Figure S2**. One walk, involving 9 PCs and 9 contigs, all with 18X ± 6X coverage, spanned about 18 Mbp total; see **Figure S2d**. These walks point to duplicated sequences, longer than individual contigs, that are present twice with low similarity, with each copy represented within a different group of adjoining contigs. Since PCs incorporate most of the assembly, the duplication most likely represents spans from homologous chromosomes that were assembled to into separate contigs.

As a third evaluation, the C6/36 contigs were subjected to BUSCO analysis [23] using genes thought to have single-copy orthologs across arthropods. Of 2624 genes found in contigs, 64% appeared as two instances. As shown in Table 1, 99.6% of two-instance genes involved 18X ± 6X contigs. There was a significant association of genes having 2 instances in the assembly and having all instances on 18X ± 6X contigs (Fisher’s exact test, p<0.01, **Table S14**). Thus, the gene duplication mirrors the redundancy observed in PCs.

**Table 1.** BUSCO gene analysis. BUSCO genes are presumed single-copy in eukaryotic genomes but most occur twice in the C6/36 assembly. BUSCO arthropod genes were searched against C6/36 contigs. Two instances were found for 1668 genes. The genes were further evaluated for whether any of their instances occurred contigs with short-read coverage in the 18X ± 6X or 36X ± 6X range. These coverage values suggest haplotype separation and collapse, respectively, within the contig sequences. Of genes with exactly two instances in the assembly, 1662 (99.6%) had at least one instance on a 18X-range contig, while only 57 had at least one instance on a 36X-range contig. This supports the characterization of 18X-range contigs as containing sequences specific to a haplotype. **Table S13** gives the coordinates and short-read coverage of every instance.

| Number of Instances | Genes Found | Genes on contigs with 18X | Genes on contigs with 36X | Genes only on contigs with 18X | Genes only on contigs with 36X | Instances on contigs with 18X | Instances on contigs with 36X | Instances on other contigs |
| --- | --- | --- | --- | --- | --- | --- | --- | --- |
| 1 | 825 | 704 | 113 | 704 | 113 | 704 | 113 | 8 |
| 2 | 1,668 | 1,662 | 57 | 1,600 | 2 | 3,262 | 59 | 15 |
| 3 | 98 | 98 | 9 | 80 | 0 | 275 | 10 | 9 |
| 4 | 24 | 24 | 1 | 23 | 0 | 94 | 2 | 0 |
| 5 to 8 | 9 | 9 | 0 | 9 | 0 | 57 | 0 | 0 |
| Total | 2,624 | 2,497 | 180 | 2,416 | 115 | 4,392 | 184 | 32 |

The situation appears nuanced for the 825 one-copy genes. Almost 14% of one-copy genes mapped to contigs with 36X ± 6X coverage, suggesting these genes are on diploid contigs represented by a consensus of two alleles. Another 1% (7 genes) hit contigs with very high (over 40X) coverage, suggesting these genes may be replicated in the genome but underrepresented in the assembly. Most single-copy genes, 85%, mapped to contigs with 18X ± 6X coverage, similar to the portion that mapped to PC contigs. Genes mapping to only one contig of a PC, suggesting a “missing gene”, were inspected further. The mapped loci did not show elevated short-read coverage, as would be expected if the gene were higher copy in the genome than the assembly. The alignments did not involve contig ends, as would be expected if the cognate gene belonged in an assembly gap, or unusual levels of discontinuity. Some “missing genes” actually did have fragmentary alignments, suggesting gene loss. As illustrated in **Table S15**, PC#1, the PC with the most shared short reads, spans 17 BUSCO genes. Its two-copy genes are ordered consistently across 6 Mbp but these are interspersed by 4 genes found on only one or the other of the two contigs, plus one gene with additional copies elsewhere in the assembly. Thus, this PC presents syntenic sequences spanning structural variants and indicates that some single-copy BUSCO genes on 18X ± 6X contigs are attributable to allele-specific gene loss.

By the three methods of read mapping, contig alignment, and gene finding, we consistently found duplication within the assembly. The results are consistent with a model of a heterozygous genome for which spans from both haplotypes are represented in the assembly. This model predicts the genome size is 52% of the assembly size, or 1.172 Gbp. Alternate models have less support. A whole-genome duplication (WGD) model predicts contig pairs. Not seen previously in mosquitoes, the WGD would have to be specific to the cell line or its ancestral strain. However, the WGD is not apparent in the C6/36 karyotypes and would not by itself predict the high intra-PC heterozygosity. Another model postulates genomic differences between the two lots of cells that were grown and sequenced separately but combined in the assembly. This model predicts the short reads would map preferentially to one contig of each PC since the short reads were derived from one lot exclusively. However, this was not the case. This model also predicts that raw long reads, mapped to contigs for consensus polish, would segregate by lot, but this was not the case (not shown). We conclude that the assembly presents both alleles at most loci. The alleles may or may not be phased, that is, contigs may not consistently derive from the same haplotype along their full lengths.

### Annotation reveals male-specific Nix, two null forms of Dicer, endogenous virus

The NCBI RefSeq annotation of contigs yielded 143,606 exons in 38,706 genes of which 28,625 are protein coding. 6,833 genes had variants and there were 42,899 mRNA transcripts. Results are public [24]. The RefSeq protein-coding gene set is 63% larger than the 17,539 protein-coding models described with the *A. albopictus* Foshan assembly [10] and likely includes allelic forms of many genes.

To assess conservation of genes and gene order between *A. albopictus* Foshan and the cell line, LiftOver [25] analysis was applied to these genomes in both directions. Of 27,093 nonoverlapping genes tested in C6/36, only 2,190 (8%) were lifted while 17,121 (63%) were split. Of 17,146 non-overlapping genes tested in Foshan, only 3,364 (20%) were lifted while 8,009 (47%) were split; **Table S16**. This analysis did not reveal high levels of gene-to-gene correspondence between the Foshan and C6/36 assemblies. This analysis may have been limited by the high dissimilarity observed between Foshan and C6/36 contigs, which would occlude context-dependent gene matching, and by consecutive alignments that hopped between homologous contigs, which would confound the recognition of conserved gene order.

The *Nix* gene in *A. aegypti* was previously shown to be located at the male-determining locus and to be necessary and sufficient to determine maleness [26]. The *A. albopictus* gene KP765684 (protein AKI28880), predicted from a partial CDS generated by transcript assembly, was established as a homolog of *Nix* based on its sequence similarity, male-specificity and transcription profile [26]. In the C6/36 assembly, KP765684 showed 100% nucleotide identity to a fragment in contig MNAF02001502 (accession NW_017857498), a 970,929 bp contig that had 17X short-read coverage indicative of allelic separation. An additional exon was identified in the C6/36 contig and it is separated by a 107 bp intron from the previously known exon that encodes KP765684. The newly predicted gene sequence encodes a 282 aa NIX protein that showed 70% similarity to the 288 aa *A. aegypti* NIX over the entire protein span. The position of the predicted intron is conserved with *A. aegypti.* Therefore, the discovery of the second exon in the C6/36 assembly extends the partial *A. albopictus Nix* gene KP765684 and further supports the homology of the *Nix* gene in the two species. The predicted protein is named “polyadenylate-binding protein 4-like protein” in the RefSeq annotation. There was no evidence of expression of this gene in our RNA sequence data from C6/36 cells at rest.

To assess the male-specificity of contig MNAF02001502, DNA sequence reads from male and female mosquitoes were mapped to the C6/36 assembly. The method of chromosomal quotient analysis [26-29] was applied to 1 Kbp spans of repeat-masked sequence. Using CQ = [(female alignments) / (male alignments)] [28] and a threshold of CQ<0.01 to indicate male specificity, fewer than 0.7% of 1 Kbp spans across the entire genome met the threshold while 14% of the 1 Kbp spans from contig MNAF02001502 did. It should be noted that the majority of the 1 Kbp spans were fully masked by repeats and did not report a CQ value. This result is again consistent with *Nix* and its contig MNAF02001502 being within the M-locus. It thus appears that the C6/36 cells are derived from one or more male mosquitoes and that they retain a full-length ortholog of NIX. These results suggest the cell line could be used to study the molecular and biochemical pathways of sex determination, including the mechanism of NIX function.

C6/36 has been observed to have a functional *dcr-1* pathway (see [30]) and a functional apoptosis pathway [31] but a dysfunctional antiviral RNA-interference response [32]. Previous genotyping of Dicer-like amplicons indicated a homozygous 1 bp deletion causing a frameshift and a premature stop codon in the C6/36 *dcr-2* gene [33]. Since the Dicer-mediated RNAi pathway has been implicated in host defense against virus in *Aedes aegypti* mosquitoes [34], the *dcr-2* null mutation suggests a mechanism for virus permissiveness in C6/36. Previously reported *dcr-2* transcript sequences align to both contigs of a PC but they have full-length alignments to one contig and a small, fragmentary alignment to the other; see **Figure S3a-b**. At the same position as the full-length alignments, C6/36 has a Dicer gene annotation marked low-quality with a note that the contig has a 1 bp deletion relative to the predicted transcript; see **Figure S3c-d**. Both contigs, and both regions, have ~19X short-read coverage. Both regions have ample support by aligned long reads; see **Figure S3e-f**. Aligned to each other, the contigs show agreement on either side of the *dcr-2* locus but not within it; see **Figure S3g-h**. These results confirm the previously reported frameshift mutation in C6/36 *dcr-2* but indicate a deletion of most of the gene in the cognate allele. The heterozygosity at this locus may have escaped notice due to a lack of matching primer sequences within the cognate allele.

Endogenous flavivirus sequences have been previously reported in C6/36 DNA [35]. Our 20X genomic short reads were mapped to the 3290 bp “Aedes albopictus containing putative integrated non-retroviral sequence” from GenBank (accession AY223844.1). The resulting coverage depth ranged from 20X to 8171X (**Figure S4**) indicating that portions of the sequence are present in the genome at high copy. The full-length viral sequence was mapped to the C6/36 assembly and found within the 5.5 Mbp contig MNAF02001791, which has 18X short read coverage. Partial matches were found at 36,476 assembly locations in 1541 contigs. A small number (7663) of RNA reads from C6/36 cells at rest aligned to the virus sequence, indicating low-level transcription. Used as a control, sequence searches of the C6/36 genome assembly did not find the densovirus C6/36 DNV, which was discovered in chronically infected C6/36 cells and appears distinct from the host genome [36].

### Transcriptomics indicates low levels of aquaporins and Kir channels

The C6/36 RefSeq transcripts were predicted using public mosquito RNA sequences without cell line RNA sequence. The RefSeq transcripts were tested for presence or absence of expression in C6/36 cells using a single run of RNA sequencing of cells at rest. Reads were mapped to transcript sequences and RPKM was computed per transcript isoform without consolidation by gene. There were 14,483 transcripts with detectable expression (RPKM ≥ 1); the mean RPKM value of these transcripts was 32 (range = 1 to 5,822). There were 1,310 highly-expressed transcripts based on a threshold of RPKM ≥ 64, i.e. having 2-fold or higher RPKM than the mean.

The highly expressed transcripts (HETs) were manually examined for the presence of genes belonging to two broad functional groups previously examined in the analysis of the *Aedes albopictus* Foshan genome [10]: detoxification proteins and odorant-binding proteins/receptors. Related to detoxification, 26 HETs were identified. Of these, 10 were cytochrome P450 oxidases (CYP450s), 2 were glutathione S-transferases (GSTs), 12 were ABC transporters, and 2 were carboxyl/cholinesterases (CCEs). Related to odorant binding proteins/receptors, 2 HETs were identified. These were orthologs of OBP9 in *An. gambiae* (AGAP000278) and OBP21 in *Ae. aegypti* (AAEL005770). See **Supplemental File “Detox_OBP”**.

HETs were subjected to a DAVID analysis (v6.7) [37, 38] to further identify functional pathways enriched among the HETs,. Among the HETs, DAVID identified 22 functional clusters that were significantly enriched (enrichment score > 1.3). By a process of manual categorization applied to mosquitoes previously [39], the enriched functional clusters were grouped into 7 broad themes: transcription and translation (8 clusters); protein sorting and trafficking(1 cluster); proteolysis (3 clusters); ATP metabolism (3 clusters); cytoskeletal functions (2 clusters); cell signaling (2 clusters); and generic (3 clusters). See Table 2 for summary and **Supplemental File “DAVID”** for details.

**Table 2.** Enrichment scores for functional clusters among the highest-expressed transcripts in C6/36 cells at rest. Functional clusters were generated and scored by DAVID and categorized manually. The ‘generic’ functional clusters contains transcripts with no specific or consistent functional theme.

| Category | Functional cluster | Score |
| --- | --- | --- |
| Transcription and Translation |  |  |
|  | Ribosome | 28.65 |
|  | Translation factor activity | 9.98 |
|  | Protein folding | 6.35 |
|  | rRNA binding | 2.26 |
|  | Elongation factor | 2.18 |
|  | Regulation of translation | 1.63 |
|  | Heat shock protein 70 | 1.58 |
|  | RNA recognition motif | 1.47 |
| Protein Sorting and Trafficking |  |  |
|  | Protein transport | 2.39 |
| Proteolysis |  |  |
|  | Proteasome | 1.62 |
|  | Ubiquitin | 1.47 |
|  | Ubiquitin mediated proteolysis | 1.31 |
| ATP Metabolism |  |  |
|  | Oxidative phosphorylation | 2.69 |
|  | Glycolysis | 2.15 |
|  | Cytochrome-c oxidase activity | 1.70 |
| Cytoskeleton |  |  |
|  | Regulation of cytoskeleton organization | 2.09 |
|  | Actin-binding | 1.39 |
| Cell Signaling |  |  |
|  | GTP-binding | 3.94 |
|  | Rho | 1.73 |
| Generic |  |  |
|  | Cellular homeostasis | 2.52 |
|  | Nucleotide binding | 2.15 |
|  | Proteasome component region | 1.46 |

This analysis suggests the C6/36 cell line is enriched with the molecular pathways for the 1) proper expression of mRNAs and proteins, 2) post-translational processing and trafficking of synthesized proteins, 3) protein turnover, and 4) synthesis of ATP. These would be expected of most cells. Similar results had been found in the transcriptome of the Malpighian tubules of nonblood fed *A. albopictus* [39]. However, the molecular pathways for 5) cytoskeletal function i.e. cell division and 6) cell signaling *i.e.* responding to environmental cues, which were not significantly enriched in the Malpighian tubules of *A. albopictus,* may reflect cell line specializations for growth in laboratory cultures.

Aquaporins (AQPs) are a family of transmembrane proteins that mediate the transport of H_2_O, small solutes (e.g., urea, glycerol), and gasses (e.g., CO_2_) across plasma membranes. Inward rectifier K^+^ (Kir) channels are a subfamily of K^+^ channels that mediate movements of K^+^ across plasma membranes. Recent work in mosquitoes has demonstrated that AQPs play key roles in water balance, heat tolerance, and vector competence, [40-45] while Kir channels play key roles in renal transepithelial K^+^ and fluid secretion and fecundity [46-50]. Kir channels are also emerging targets for mosquitocide development [46-48, 51]. Typically, the genomes of mosquitoes possess 6 genes encoding aquaporins: Drip (AQP1), Prip (AQP2), Bib (AQP3), Eglp1 (AQP4), Eglp2 (AQP5), and Aqp12L (AQP6); and at least 4 discrete genes encoding Kir channels: Kir1, Kir2A, Kir2B, and Kir3. There is some gene duplication in the C6/36 assembly, for which the 25 annotated AQP protein isoforms can be assigned to 15 haploid alleles from 9 (possibly 8) diploid loci, based contig coordinates and PC relationships. Likewise, the 15 annotated Kir protein isoforms indicate some gene duplication of the 4 genes expected. Despite the robust number of AQP and Kir genes in C6/36 cells, AQP mRNA expression was not detected via RNA sequencing *(i.e.* RPKM < 1), and only two isoforms of Kir1 were nominally expressed (i.e. RPKM = 1), suggesting that the cell line is deficient in AQP-mediated water, solute, and gas transport, and Kir-mediated K^+^ transport. See **Supplemental File “AQP_Kir”**.

### Demonstration of subtraction database for viral assays

Host subtraction is the bioinformatics process of filtering reads derived from host DNA and RNA, thereby enriching non-host reads [52]. Host subtraction assists the discovery and characterization of viral sequences present at low titer in voluminous short-read datasets [52]. To evaluate utility for host subtraction in viral assays, the C6/36 transcript and genome sequences were used to filter RNA sequence reads from 26 samples of Zika-infected and 6 samples of mock-infected C6/36 cells. Quantitative PCR confirmed Zika RNA was abundant in the Zika-infected samples vs. the mock-infected samples. A single multiplex sequencing library was prepared by the low-cost SISPA method [53]. Unpaired Illumina RNA sequencing reads were filtered by mapping to C6/36; see **Table S17**. Transcript mapping removed 13.2%, and genome mapping further removed 76.1%, of total reads. The remaining 10.7% of reads were given a taxon assignment by blastn best hit to the NCBI non-redundant nucleotide database. Of total reads, 1.10% received a taxon assignment including 0.011% assigned to Zika, and zero to any other viral taxon, indicating that Zika was detected accurately by this assay. Critically, subtraction reduced cpu time to 15% of what would have been required to blast all reads. Subtraction using instead the *Ae. albopictus* Foshan sequences was slightly less effective leaving 13.2% of reads instead of 10.7% to be characterized by blast.

## DISCUSSION

The C6/36 genome assembly offers large contigs attributable to deep coverage by long-read sequencing. Longer than the contigs of any previously assembled mosquito genome, the large contigs offer not only complete gene sequences but also gene context including repetitive DNA. The contigs are not joined by scaffolds and they are not mapped to chromosomes, though physical mapping technologies including Hi-C, Dovetail, and BioNano technologies (reviewed in [54]), would be able to make use of our contigs. The contigs are publically available with gene annotation provided by the NCBI Eukaryotic Genome Annotation Pipeline. We believe C6/36 becomes the second cell line, after HeLa [55], to have its genome *de novo* assembled. The accuracy of the assembly is supported by several observations. Short-read data that had not been used during assembly aligned to the assembly at a high rate with high identity. Alternate assemblies of the long reads, including those generated using different software, were very similar as shown by local alignment and short-read mapping.

Most of the assembly contains duplicated sequence though our karyotype analysis did not indicate a whole-genome duplication. We demonstrated that the duplication captures haplotype variants of the diploid genome. The duplication complicates analysis by gene count but it makes the assembly a valuable reference for read mapping and for detection of allelism. Some recent genome projects intentionally separated homologous sequences during assembly [17, 56] but the separation within the C6/36 assembly was a byproduct of heterozygosity within the genome. With additional resources, it might be possible to identify haplotype-phased blocks within contigs or to organize contigs into haplotype-phased scaffolds, *e.g.* [57].

If the C6/36 assembly were complete and fully haplotype-separated, the total span would be twice the genome size. Compared to the *Aedes albopictus* Foshan mosquito assembly [10], the C6/36 contig span is only 28% larger. Some degree of haplotype separation may be present in the Foshan assembly and the two assemblies may represent different strains or genomes of different size. Alignments with at least 95% identity covered small portions of both assemblies.

In the C6/36 assembly, we discovered a two-exon sequence for *Nix,* the male-specific gene in *Aedes albopictus.* We confirmed maleness of the cell line through differential mapping of reads from male and female mosquitoes as well as by the identification of the M factor *Nix.* This finding could be helpful for testing sex-specific agents developed for mosquito sterility programs. We also discovered that the cell line’s *dcr-2* locus, source of the Dicer homolog in mosquitoes, contains a second null mutation allelic to the previously described truncated form.

Using RNA sequencing of cells at rest, and the RefSeq annotation of the C6/36 assembly, we noted the conspicuous absence of mRNAs encoding AQPs and Kir channels. Our analysis of AQPs and Kir channels suggested the cell line is deficient in AQP-mediated water, solute, and gas transport, as well as Kir-mediated K^+^ transport. It is intriguing to speculate that the nominal AQP and Kir mRNA expression is an adaptation of the cell line to stable cell culture conditions wherein the extracellular environment, *i.e.* culture media, is not subject to fluctuations in osmolality, K^+^, or temperature as would be experienced on a regular basis in the mosquito. Given that Kir channels in *Drosophila melanogaster* have been implicated in the RNA interference antiviral immune pathway [58], it is also possible that the nominal Kir mRNA expression contributes to the susceptibility of C6/36 cells to arboviral infection. It is unlikely that the original mosquito cells that generated the cell line would be deficient in AQP and Kir mRNA expression given the near ubiquitous expression of at least one AQP and Kir mRNA in various mosquito tissues that have been previously examined [39-45, 50, 59-62]. The lack of endogenous AQP and Kir mRNA expression may be serendipitous as it suggests that C6/36 cells have potential to offer a mosquito-based cell line for functionally-characterizing mosquito AQPs and Kir channels, if the cells can be transfected with and induced to express exogenous cDNAs. The genome sequence should enable more extensive transcriptomics including of cells at various stages of viral infection.

## POTENTIAL IMPLICATIONS

Our results should enable further use of the C6/36 cell line for virus detection, virus surveillance, vaccine and antiviral drug development, and promoting a basic understanding of the virus-host interplay for medically important mosquito-transmitted viruses. The genome sequence can be used as a bioinformatics filter to remove host sequence and thereby enrich nonhost reads among DNA or RNA sequence data from exposed cells. Using the assembly as a filter could avoid uncertainty that would be caused by the endogenous viral sequences whose presence we confirmed in the genome and the transcriptome. Filtering with a largely-complete genome sequence will ease detection of novel and low-titer viruses. The genome sequence may enable discovery of microRNAs expressed by the cells in response to specific conditions. The genome sequence and annotation will enable reference-guided expression studies that characterize and quantify viral progression and host response. Applications for the cell line could expand if transcriptome studies reveal active pathways that could be targeted by insecticides or inactive pathways that could be studied by ectopic expression of insect genes.

## METHODS

### Sequencing

Cells were obtained from two independent shipments of *Aedes albopictus* clone C6/36, ATCC CRL-1660 from ATCC (Manassas VA). Cells were maintained in Minimal Essential Media supplemented with 10% fetal bovine serum and nonessential amino acids. Cells were maintained at 28°C and 5% CO2, and confluent monolayers were harvested by cell scraping. Following thaw, cells were passaged a single time and subjected to genomic DNA isolation (Qiagen, Germantown MD). DNA for PacBio sequencing was subjected to library construction following manufacturer instructions [63]. DNA from lot #59479117 was sequenced at NBACC (Fort Detrick, MD) using 128 SMRT cells and multiple libraries. DNA from lot #62871143 was sequenced at Icahn School of Medicine at Mt. Sinai (NY) using 80 SMRT cells. All sequencing used PacBio RS II instruments with P6C4 chemistry. Raw reads were extracted as subread FASTQ files from instrument h5 files using SMRTlink software. Lot #62871143 was also used for Illumina sequencing. Genomic DNA was Blue Pippin (Sage Science, Beverly MA) sheared to generate 270 bp fragments. These were end-repaired, A-tailed, and ligated to Illumina adaptors following standard protocols (NEB, Ipswich MA). Libraries were subjected to Illumina NextSeq 2x150 bp paired end (PE) sequencing. Total RNA from C6/36 cells at rest was isolated with RNAeasy (Qiagen). Total RNA from cells at rest was enriched for messenger RNA with oligo dT dynabeads (Invitrogen, Carlsbad CA). Total RNA from mock-infected and Zika-infected samples were subjected to SISPA multiplex library construction [53] and sequenced by Illumina NextSeq 1x150 bp sequencing.

### Genome assembly

All Falcon [17] assemblies were generated on the DNAnexus (San Francisco CA) platform using the DNAnexus Falcon 0.0.1 application which combined Falcon 0.4.2 with REPMask and TANMask from DAMASKER [64]. Raw reads of 10,731 bases or longer were subject to error correction. Corrected reads of 10,000 bases or more were subject to overlap and contig computation. The Falcon p_contig and a_contig sets were combined for analysis. All Canu [19] assemblies were generated with Canu [65] with the command ‘canu errorRate=0.013 ‐p asm ‐d C636_canu genomeSize=2g’. Falcon and Canu contigs were subject to consensus polishing using all the raw PacBio reads and either one or two iterations of Quiver or Arrow from PacBio SMRTlink. Falcon polishes used SMRTlink 3.1 on the DNAnexus platform. Canu polishes used SMRTlink 3.1.1 on an SGE grid; see [66]. The N50 statistic shows the length of the shortest contig such that contigs of equal or greater length span at least half of some total, usually the assembly size. The NG50 uses the putative genome size, **G**, and for our NG50 calculations, **G** was set to the total span of the contigs from Canu and 2 rounds of Arrow (2,247,306,400 bp). Two contigs, each composed of a homopolymer repeat, were removed from the final assembly.

### Repeats, maps, alignments

Repeats were detected with the Repeat Modeler package [67-71] version 1.0.8. Short-read K-mer analysis was generated with GenomeScope [72] version 1.0 with k=21. Short reads were mapped to assemblies with bowtie2 version 2.2.5 [73]. Mappings were restricted to concordantly mapped pairs under default parameters corresponding to end-to-end alignment and sensitive settings for one best mapping per pair with ties broken randomly, with the following exceptions. C6/36 genomic short-reads were mapped to the Foshan assembly in “very sensitive” mode. For detection of Paired Contigs (PCs), short reads were mapped to C6/36 contigs with the “best 4” parameter. The mapping of RNA reads to virus sequence was filtered for MapQ≥5. Repeats were mapped to C6/36 contigs in local sensitive mode retaining all alignments. Outputs were analyzed in bam format with samtools and bedtools [74, 75]. Mode contig coverage is computed as the center value of a 3 wide window starting at 3 fold coverage (3X) having the most bases in that coverage window; for example, the minimum coverage window, 3-5X, is reported as 4X. Contig local alignments were generated with nucmer 3.1, part of the MUMmer package [76] compiled for 64-bit processors and filtered with delta-filter -1, minimum 1000 bp. Local alignment dot plots were visualized with mummerplot. Global alignments were generated with ATAC [77] which computes maximal chains from 1-to-1 K-mers. Aligned spans were accumulated over stringent and permissive chains, denoted ‘M r’ and ‘M c’, respectively, in the outputs.

### Gene annotation and analysis

Single-copy genes were downloaded from BUSCO v1.22 arthropoda-odp9 [23]. *Nix* was mapped with blast [78]. The contig sequence was annotated by the NCBI Eukaryotic Genome Annotation Pipeline 7.2 [79]. Evidence included alignments of 5.6 G public *Aedes albopictus* RNA sequences (excluding C6/36 RNA sequence generated by this project) and 137 K public insect protein sequences. The annotation pipeline was adjusted to accommodate long introns containing TE-associated ORFs as in *Aedes aegypti* [13], after such introns were detected in an initial run on C6/36. Mapping of gene annotations between assemblies was performed with LiftOver [25] which uses chained BLAT [80] alignments to transfer coordinates. Chromosomal quotient analysis to identify male-specific contigs [26, 28] used C6/36 contigs repeat masked [81] and split into 1 Kbp spans; reads were mapped by bowtie [82] with parameters ‐a ‐v 0. Pathway enrichment analysis was performed using DAVID (v6.7) [37, 38] and best blastp hits in *An. gambiae* or *Ae. aegypi.*

### Transcriptome analysis

Mosquito proteins were taken from VectorBase release VB– 2016-12. Best blast hits were found with NCBI blastall 2.2.26 [78] using default parameters and tabular outputs. Genes related to diapause etc. were taken from [10] supplemental documents SD9, SD13, SD15., and translated from *Aedes aegypti* Liverpool (AAEL) accession to the best blastp hit in C6/36.

### Growing cells for karyotype

Cells were grown in DMEM (Gibco, 11995065) supplemented with non-essential amino acids (Gibco, 11140050) and L-glutamine (Gibco, 25030081) at 28°C supplemented with 5% CO2. Once cells were 80% confluent, approximately 10^5^ cells were passaged into a 6 well plate by scraping. Once cells had settled and reattached to the plate, fresh C6/36 media was supplemented with Colcemid (ThermoFisher, 15212012) so that the final concentration was (1 μg/ml). After a 2 h incubation at 28°C with 5% CO2 media was aspirated and cells rinsed with PBS (Gibco, 14040133). Cells were scraped and resuspended in PBS.

### Fixing cells for karyotype

The cells suspended in PBS were centrifuged at 1000 rpm for 4 minutes, aspirated supernatant, leaving about 200 μl in the tube. The bottom of the tube was tapped to break any clumps and added 5 ml of ice cold 0.56% KCl solution, inverted once. The cells were incubated at room temperature for 6 minutes and centrifuged at 1000 rpm for 4 minutes. The supernatant was aspirated, leaving 50 μl in the tube. The pellet was resuspended by gently tapping the bottom of tube. The cells were fixed by adding 5 ml of methanol: glacial acetic acid (3:1) fixative solution, the fixative was added one or two drops at a time for the first 4 ml and tapping the bottom of the tube to mix cells continuously. The fixed cells were centrifuged at 1000 rpm for 4 minutes, aspirated the supernatant and suspended the cells pellet in 200 μl of fixative.

### Visualizing chromosomes for karyotype

The fixed cells suspension (approx. 10-50 μl) was put on an alcohol cleaned slide and air dried for 1 hour. Then 10 μl of 30 nM DAPI (Invitrogen, D1306) solution in PBS was added to each slide. The slide was coverslipped and incubated in dark at room temperature for greater than 20 minutes. The coverslip was removed and slide was rinsed thoroughly with PBS. The slides were visualized on a Axioskop 2 plus (Zeiss, Oberkochen, Germany) fluorescence microscope using DAPI filter, under oil immersion at 1000X magnification. Images were taken with AxioCam MRc5 (Zeiss) camera using AxioVision software. The color images were converted to grayscale, inverted, cropped, and adjusted for brightness and contrast in Photoshop (Adobe Systems, San Jose CA).

## DECLARATIONS

### List of abbreviations

AQP: aquaporin gene
ATCC: American Type Culture Collection, provider of C6/36 cells
BUSCO: Benchmarking Universal Single-Copy Orthologs, analysis of genes thought to be unique in Eukaryotic genomes
C6/36: A cell line derived from *Aedes albopictus.*
Canu: A genome assembler derived from Celera Assembler
CQ: the Chromosomal Quotient used to measure male-specificity
HET: a Highly Expressed Transcript
K-mer: here, a string of consecutive nucleotides with a specific length, K
PacBio: Pacific Biosystems, its sequencing platform, or the read type it generates
N50: the length of the shortest member in the smallest set of contigs required to span 50% of the assembly size
PC: as defined here, Paired Contigs that contain similar sequences
RPKM: RNA read count normalized by Reads Per Kilobase of transcript per Million reads

### Funding

JCVI staff was supported by DHS contract HSHQDC-15-C-B0059. XGC was supported by the National Nature Science Foundation of China (81420108024) and the Natural ScienceFoundation of Guangdong Province (2014A030312016). ZT was supported by NIAID grant AI123338. PMP was supported by state and federal funds appropriated to the OARDC of the Ohio State University. SK and AMP were supported by the Intramural Research Program of the National Human Genome Research Institute, National Institutes of Health. NHB and TS were supported under Contract No. HSHQDC-07-C-00020 awarded by the Department of Homeland Security (DHS) Science and Technology Directorate (S&T) for the management and operation of the National Biodefense Analysis and Countermeasures Center (NBACC), a Federally Funded Research and Development Center. The views and conclusions contained in this document are those of the authors and should not be interpreted as necessarily representing the official policies, either expressed or implied, of the DHS or S&T. In no event shall the DHS, NBACC, S&T or Battelle National Biodefense Institute (BNBI) have any responsibility or liability for any use, misuse, inability to use, or reliance upon the information contained herein. DHS does not endorse any products or commercial services mentioned in this publication.

### Authors’ contributions

Manuscript: JRM, PMP, RSS, ZT, IVS, MVS. Sample prep: KD. Karyotype: IVS, MVS, DMB, VP. Sequencing: RS. Assembly & sequence analysis: SK, AMP, JRM, GS. Repeats: DH. Liftover: BR. Annotation: FTN. Nix: ZT, XGC. Dicer: JRM. Flavivirus: RSS. Expression: PMP. Subtraction: JRM. Project conception: RS, TS, NHB, GGS, AMP, RSS.

## Acknowledgements

We are grateful for contributions from Chai Fungtammasan and Brett Hannigan at DNAnexus; Karen Beeri formerly at JCVI; Carlos J. Esquivel at OSU; Yang Wu at VT.

